# A Manifold Framework for Interpretable Brain Age Estimation and Aging Trajectory Mapping

**DOI:** 10.1101/2025.09.24.678174

**Authors:** Zifei Liang, Collin J. Szczepanski, Chenyang Li, Jiayi Li, Henry Rusinek, Yulin Ge, Jiangyang Zhang

**Author notes:** Corresponding to: Jiangyang Zhang. These authors contributed equally to this work (Zifei Liang, Collin J. Szczepanski).

## Abstract

Brain aging involves complex structural changes that challenge traditional analytical methods and hinder personalized assessment of brain health. Integrating regional brain alterations into a unified, interpretable representation is difficult due to the high dimensionality of neuroimaging data. Here, we projected regional brain volumes from the Human Connectome Project Aging Dataset onto a low-dimensional manifold that reflects underlying neuroanatomical constraints. We then built a transparent framework on the manifold to estimate brain age and identify key regional drivers of the estimate. By analyzing local neighborhoods on the manifold, we identified distinct structural aging trajectories, including pronounced frontal atrophy occurring predominantly in males. This approach provides a biologically interpretable means of characterizing individual brain-aging patterns, reveals heterogeneous aging pathways, and supports more personalized assessments and insights into the aging process.

**Teaser:** A manifold representation can map brain aging patterns across regions and reveal distinct trajectories

## Introduction

Brain aging is a highly heterogeneous process, marked by varying rates of changes across age, brain regions, and individuals. This variability extends to neurodegenerative diseases, such as Alzheimer’s disease (AD) and AD-related dementia, which disrupts the course of normal aging in region-specific manners (*1–3*). Although magnetic resonance imaging (MRI) studies have consistently identified robust age-related alterations in multiple brain regions (e.g., ventricular enlargement (*4*) and hippocampal atrophy(*5*)), the spatiotemporal heterogeneity of aging processes leads to complex regional patterns that vary substantially between individuals (*6*, *7*). It poses significant challenges for providing precise personalized assessments of brain and cognitive health, as well as for making accurate diagnoses or prognoses along the continuum from normal to pathological aging.

To address this challenge, researchers have utilized large-scale MRI datasets and machine learning algorithms to estimate brain age (*8*)—the biological age of the brain—based on whole-brain MRI scans. The brain age gap, defined as the difference between predicted brain age and chronological age, has shown stronger associations with cognitive performance and health outcomes than chronological age alone, with typical prediction accuracies of mean absolute error (MAE) around 3-7 years in healthy adults(*9–11*). However, the brain age metric lacks biological interpretability and offers little insight into the specific regional changes underlying the overall assessment. Even though normative trajectory models have been developed to track multiple brain regions across the lifespan, offering standard regional aging curves along with individual centile scores(*12–16*), capturing the complexity of multi-regional dynamics remains difficult due to the large number of brain regions, inter-regional correlations (*17*), and substantial inter-individual variability. In response, data-driven approaches have begun to identify distinct clusters of brain regions with interconnected aging dynamics, offering new ways to examine the aging process (*18*, *19*). Together, these advances pave the way for developing a new metric that integrates intuitive global brain age with personalized regional aging profiles, thereby enhancing both clinical utility and biological transparency.

Notably, while MRI data are inherently high-dimensional, aging-related changes in the brain are likely confined to a lower-dimensional manifold (*20*). This concept, known as the manifold hypothesis (*21*), suggests that both inter-individual variations and age-related changes are governed by underlying anatomical constraints, rather than occupying the full high-dimensional space. Previous studies have demonstrated the existence of such manifolds and used them to distinguish AD from normal aging (*22*, *23*). However, the development of interpretable brain age metrics that utilize these compact manifold representations remains unexplored.

In this study, we built a manifold representation of the brain aging continuum based on structural MRI data from the Human Connectome Project Aging (HCP-A) dataset (*24*) and developed an associated computational framework. Specifically, we characterized age-related anatomical changes by analyzing data within neighborhoods on the manifold, where subjects exhibited similar regional volume configurations. We then integrated findings from these anatomically homogeneous subgroups across the entire manifold to reveal large-scale patterns of brain aging. Unlike conventional deep learning–based brain age models, which often operate as opaque systems (*25*), this framework enabled us to provide overall brain age estimates and identify the specific anatomical changes that most strongly influence these estimates, offering biological interpretability at the regional level.

## Results

### A two-dimensional (2D) representation of the aging brain continuum

T_1_-weighted MRI data from 690 HCP-A subjects were parcellated into 267 regions using an automated multi-atlas pipeline (**Fig. 1a**), producing a 267×1 regional volume vector V for each subject (**Fig. 1b**). We utilized uniform manifold approximation and projection (UMAP)(*26*) to construct a mapping <. that projects all subjects’ *v*s onto a 2D manifold, called aging manifold here (**Fig. 1c**). Manifold stability was assessed by systematically varying UMAP parameters (**Supplementary Materials Fig. S1**). The distribution of HCP-A subjects on the manifold showed gradients corresponding to age (**Fig. 1d**) and sex (**Fig. 1e**), consistent with well-established sex and age-related differences in brain morphology. Similar gradients were also found in results from principal component analysis (PCA) (**Supplementary Materials Fig. S2a-b**), although the subject distribution was less uniform, likely because that PCA was driven primarily by the large differences between young and old subjects (*27*). Normalizing V by individual intracranial volume removed the sex-related gradient while preserving the age-related gradient (**Supplementary Materials Fig. S3a-b**). Several outliers that deviated from the age-related gradient were observed (**Fig. 1d**); however, the overall brain morphology of these outliers aligned with their manifold neighbors better than age-matched subjects, suggesting mismatches between chronological ages and expected regional volumes. By stratifying male and female HCP-A subjects into multiple age groups, we computed average aging trajectories for each sex based on mean regional volumes and their 95% confidence intervals on the manifold (**Fig. 1f**). The substantial scatter of individual subjects around these average trajectories highlighted the considerable heterogeneity of aging processes.

**Figure 1:**
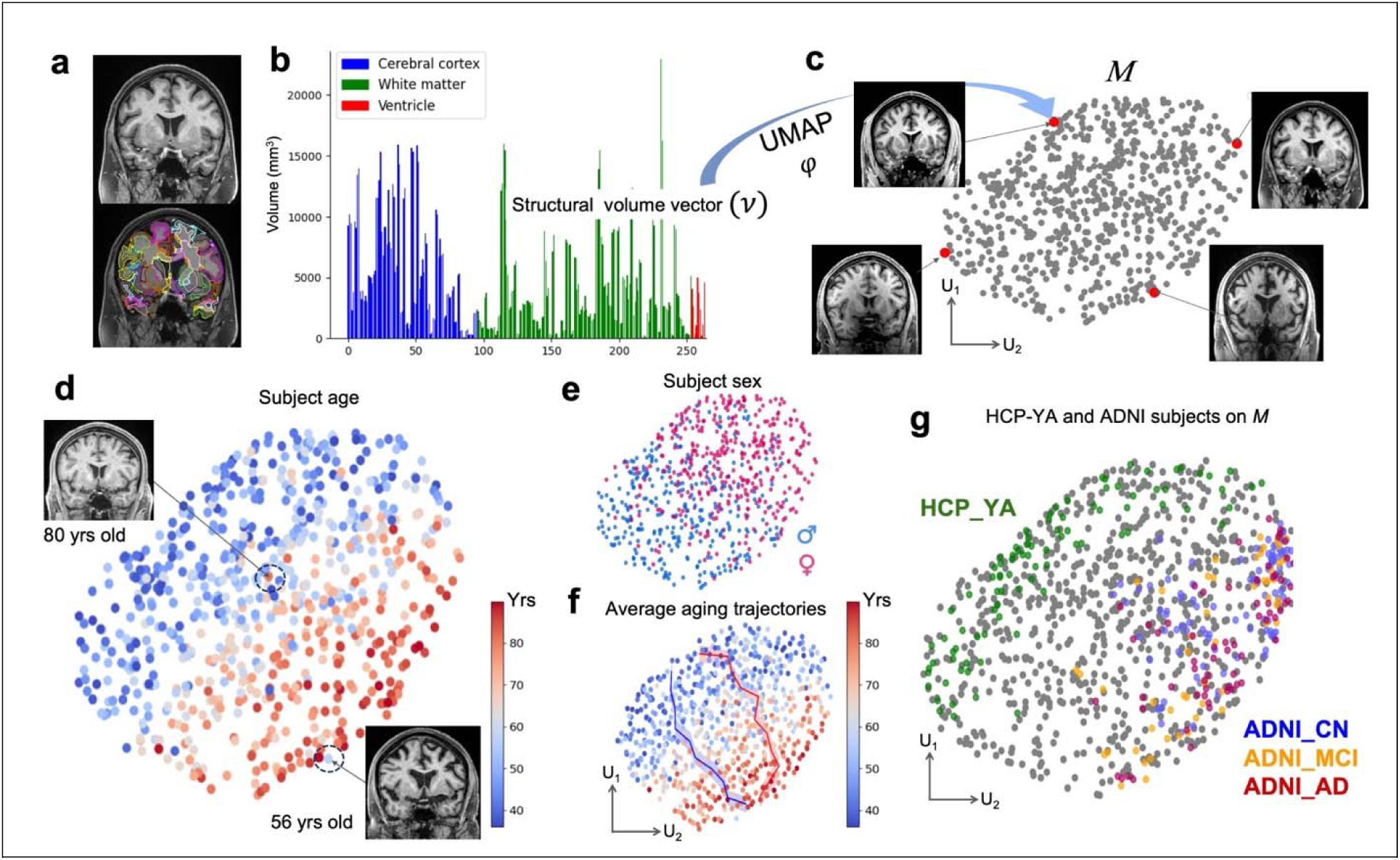
Mapping of high-dimensional regional volume data from the HCP-A dataset onto a 2D manifold using UMAP. **a-b:** T_1_-weighted MRI images from each HCP-A subject were segmented into 267 regions, whose volumes formed a volume vector V. **c:** The volume vectors of 690 HCP-A subjects were projected using UMAP onto a 2D manifold M, visualized as a point cloud where each point represents one subject. **d**: Distribution of chronological age across individual subjects on M. The insets show two selected outliers whose chronological ages differed markedly from their neighbors’ but who shared similar brain morphology (e.g., ventricle size) as shown in panel **c**. **f:** Average aging trajectories of male and female subjects on M with 95% confidence intervals. **g:** Distribution of external validation datasets on M: HCP-YA subjects (green) and ADNI subjects, color-coded by diagnostic status (red for AD, orange for mild cognitive impairment (MCI), and blue for cognitively normal (CN)).

The aging manifold, representing plausible regional volume configurations along the aging continuum, can be generalized to other datasets. Although the manifold was constructed using the HCP-A dataset, structural MRI data from other studies can also be mapped onto it using the UMAP-derived <.. To test this generalizability, we mapped selected data from the HCP-Young adult (HCP-YA)(*28*) and Alzheimer’s disease neuroimaging initiative (ADNI)(*29*) datasets onto the manifold after harmonization (**Fig. 1g**). The HCP-YA subjects (ages 25-40 years) colocalized with younger HCP-A subjects, while the ADNI subjects (ages 55-90 years) positioned alongside older subjects at the opposite end of the manifold. Similar but more scattered results were observed in the PCA-based analysis (**Supplementary Materials Fig. S2c**). The aging manifold can be readily extended by incorporating structural MRI data from additional datasets (e.g., HCP-YA, as shown in **Supplementary Materials Fig. S3c**) or even non-MRI data modalities (e.g., cognitive test results, as demonstrated in **Supplementary Materials Fig. S3d-e**).

### Estimation of brain ages on the manifold

The observed age-related gradient on the manifold suggested that age could be modeled as a smooth function across the manifold, enabling estimation of a subject’s brain age. To achieve this, we used Watson kernel regression within local neighborhoods on the manifold, where each neighborhood consisted of subjects with similar regional volume profiles and known chronological ages (see inset in **Fig. 2a**). We determined that a neighborhood size of approximately 50 subjects provided optimal performance. Using leave-one-out validation, we found a strong correlation between estimated and chronological age (R^2^=0.74, p<0.0001, **Fig. 2a**), with a MAE of 5.8 years. This accuracy was comparable to previously published deep learning models trained on similar sample sizes (*11*), though it was higher than MAEs reported for models trained on much larger datasets (10,000 or more subjects)(*30*, *31*). Brain age estimates derived from a 3D manifold showed similar performance (R^2^=0.73, p<0.0001, MAE = 6.1 years).

**Figure 2:**
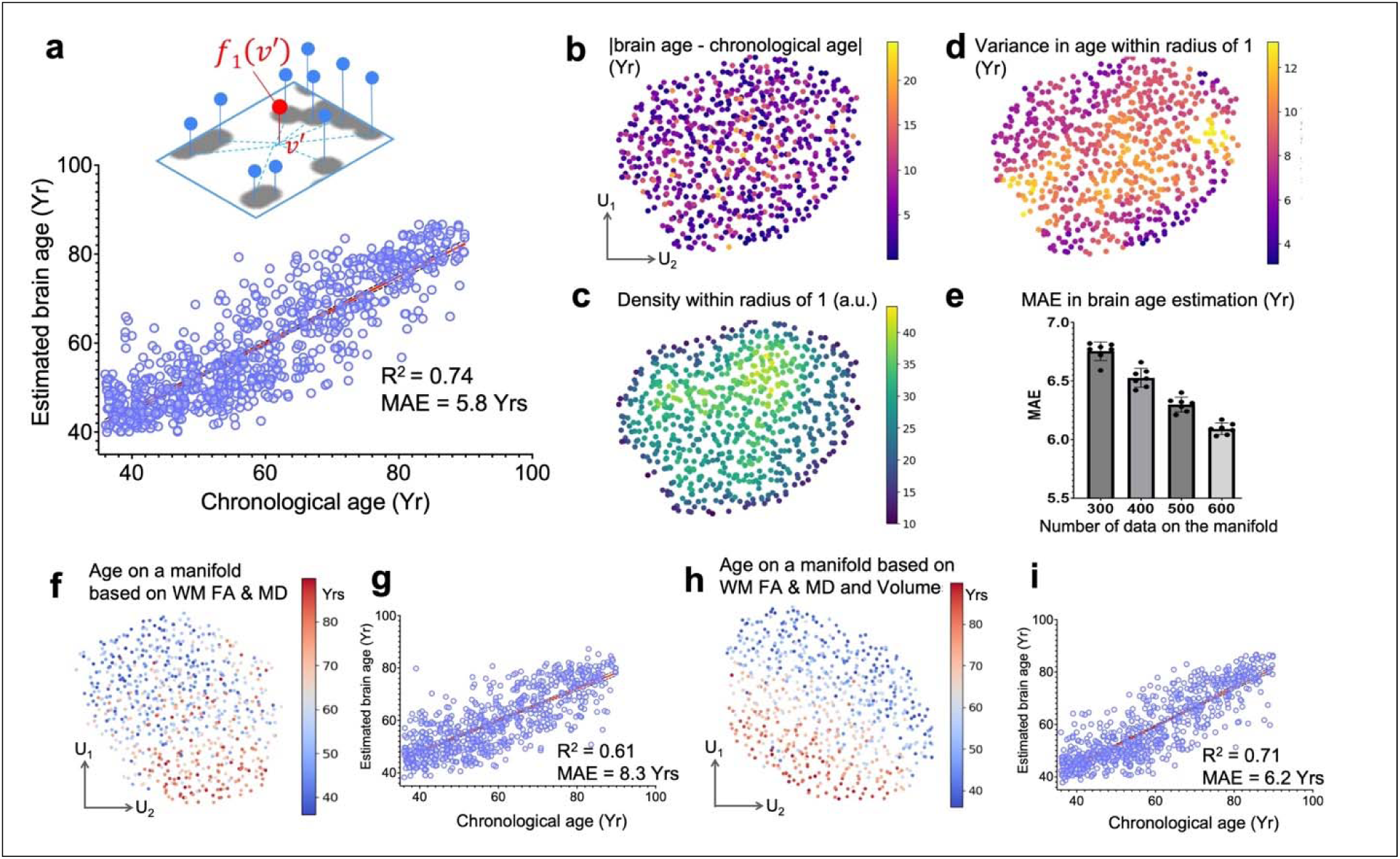
Brain age estimation on the aging manifold. **a:** Correlation between estimated brain age and chronological age for HCP-A subjects. The inset shows a schematic of regression for a subject with a regional volume vector (***v’***, red) based on existing data points (blue) within local coordinates corresponding to a neighborhood on the manifold. **b:** Absolute differences between the estimated brain age and chronological age across the manifold. **c:** Number of subjects within a unit sphere for each point on the manifold (local density). **d:** Variance of chronological ages within each subject’s neighborhood. **e:** MAEs of brain age estimation as a function of the number of subjects included on the manifold. **f:** 2D manifold constructed using white matter fractional anisotropy (FA) and mean diffusivity (MD). **g:** Brain age estimation based on white matter FA and MD showing strong correlation with chronological age. **h-i:** 2D manifold constructed using both regional volumes and white matter FA/MD (**h**), and corresponding brain age estimation performance (**i**).

The estimation error was distributed relatively uniformly across the manifold (**Fig. 2b**) and was influenced by the density of subjects and the variation in ages within each neighborhood (**Fig. 2c-d**). For example, when using a randomly selected subset of 600 subjects, the average MAE of brain age estimation increased to 6.1 years and continued to increase as subject density was further reduced (**Fig. 2e**). Furthermore, brain age estimation for 200 randomly selected HCP-YA subjects yielded an MAE of 10.2 years, likely because HCP-YA subject ages fell outside the age range of the HCP-A training dataset. However, using an extended manifold constructed from both HCP-A and HCP-YA datasets (**Supplementary Materials Fig. S3c**), the MAE for brain age estimation of an independent set of HCP-YA subjects was reduced to 5.5 years.

The method can be extended to incorporate additional MRI markers. We constructed a 2D manifold using fractional anisotropy (FA) and mean diffusivity (MD) data—two widely used diffusion MRI markers—from white matter regions (**Fig. 2f**). Brain age estimation based on white matter FA and MD from 600 subjects yielded an MAE of 8.3 years and R² of 0.61 (**Fig. 2g**). Combining regional volumes and white matter FA and MD values (**Fig. 2h-i**) improved MAE to 6.2 years (R² of 0.71 with 600 subjects on the manifold).

### The manifold encoded regional volume information

Whole brain volume varied gradually across the manifold (**Supplementary Materials Fig. S4a**), indicating that subjects with similar volumes clustered together. Similar to age, regional volume information was encoded by positions on the manifold. Leave-one-out validation revealed a strong correlation between estimated and actual whole brain volumes (**Supplementary Materials Fig. S4b**, R^2^=0.99). Individual structures showed lower but still strong correlations (**Supplementary Materials Fig. S4c-g**), potentially due to the regional volumes being noisier than whole brain volume. A 3D manifold further improved correlation strength and volume estimation accuracy (**Supplementary Materials Fig. S5**), suggesting it captured additional structural information beyond the 2D representation.

### Regional volumes that contribute to brain age estimation

Understanding how individual regional volumes contribute to the estimated brain age is crucial for biological interpretability. The smooth gradients observed across the manifold (**Fig. 1d & Supplementary Materials Fig. S4**) suggested that subjects within each neighborhood shared similar structural volumes and chronological ages, forming a relatively homogeneous subgroup of the HCP-A dataset. This local homogeneity provided the necessary conditions for approximating the relationship between regional volumes and age using a multivariable linear function within each neighborhood. To address collinearity among structural volumes (**Supplementary Materials Fig. S6**), we employed partial least squares regression (PLSR) to estimate local relationships and ranked input markers based on variable importance in projection (VIP) scores (**Fig. 3a**). Seventy-two regions consistently maintained VIP scores above 1 across the manifold (**Fig. 3b-d**), where VIP > 1 is typically considered indicative of high impact. Among these, frontal sulcus and cerebellar gray matter exhibited the highest average VIP scores, followed by parietal sulcus, frontal lateral ventricles, superior temporal gyrus, and medial frontal gyrus (**Fig. 3e-h**). Brain age estimation using a new manifold constructed from these 72 key structures achieved an MAE of 6.1 years (**Supplementary Materials Fig. S7a-b**), matching the performance of all 267 structures and confirming that these regions contained equivalent information for brain age estimation. Incorporating cognitive function data further reduced the MAE to 5.9 years (**Supplementary Materials Fig. S7c-d**).

**Figure 3:**
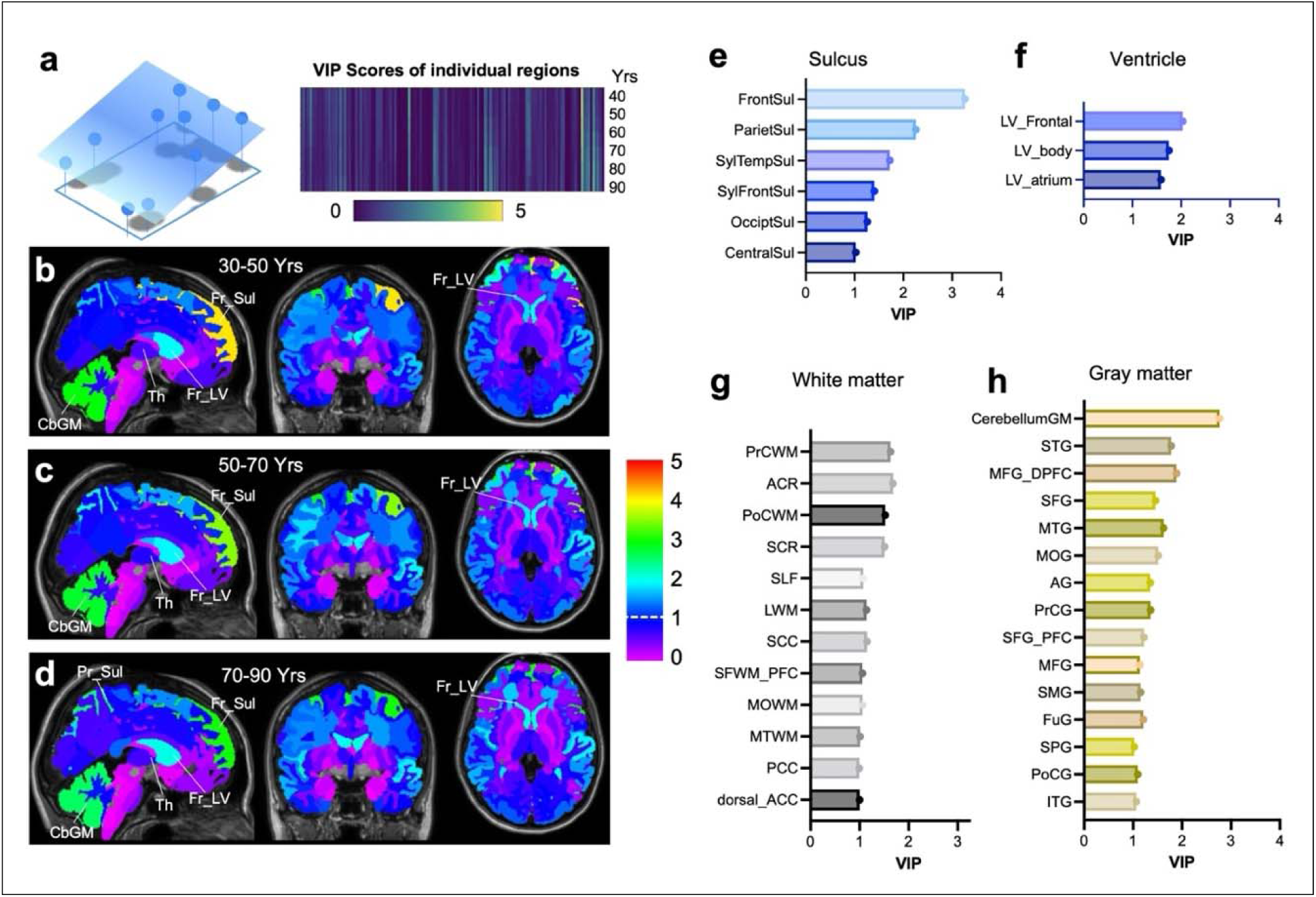
Local analysis of individual brain regions’ contributions to brain age estimation. **a:** Schematic illustrating how the brain age function is locally approximated using multivariable linear functions, from which VIP scores are calculated for each brain structure. **b-d:** Maps of regional VIP scores for subjects in three age groups: 30-50 years, 50-70 years, and 70-90 years. **e-h:** Average VIP scores of the 72 high-impact structures organized into four anatomical categories, with scores shown for both left and right hemispheres. Brain region abbreviations are provided in **Supplementary Table S1**.

VIP scores of individual regions varied with age, reflecting their changing contributions to brain age estimation. The frontal lateral ventricle (LV) VIP score increased with age (**Fig. 4a**), consistent with its relatively stable volume in subjects aged 40-50 years followed by accelerated expansion at later ages (**Fig. 4b**). The frontal LV volume increased more rapidly in males than females, explaining males’ higher VIP scores for this region. In contrast, two white matter structures—superior corona radiata (SCR) and anterior corona radiata (ACR)—showed steady age-related volume declines with relatively constant VIP scores. Interestingly, the thalamus maintained a VIP score near one despite significant age-related changes, likely because its volume was highly correlated with other brain regions, including ACR and SCR (**Supplementary Materials Fig. S8**). High correlations between regional volumes reduce individual VIP scores because correlated regions provide redundant rather than unique information for age prediction. For example, hippocampal volume had a lower VIP score due to strong correlations with middle temporal gyrus (MTG), ACR, and SCR volumes. A separate PLSR analysis revealed that 99% of hippocampal volume variation could be predicted from the 72 key structures, rendering it redundant for age estimation in this framework.

**Figure 4:**
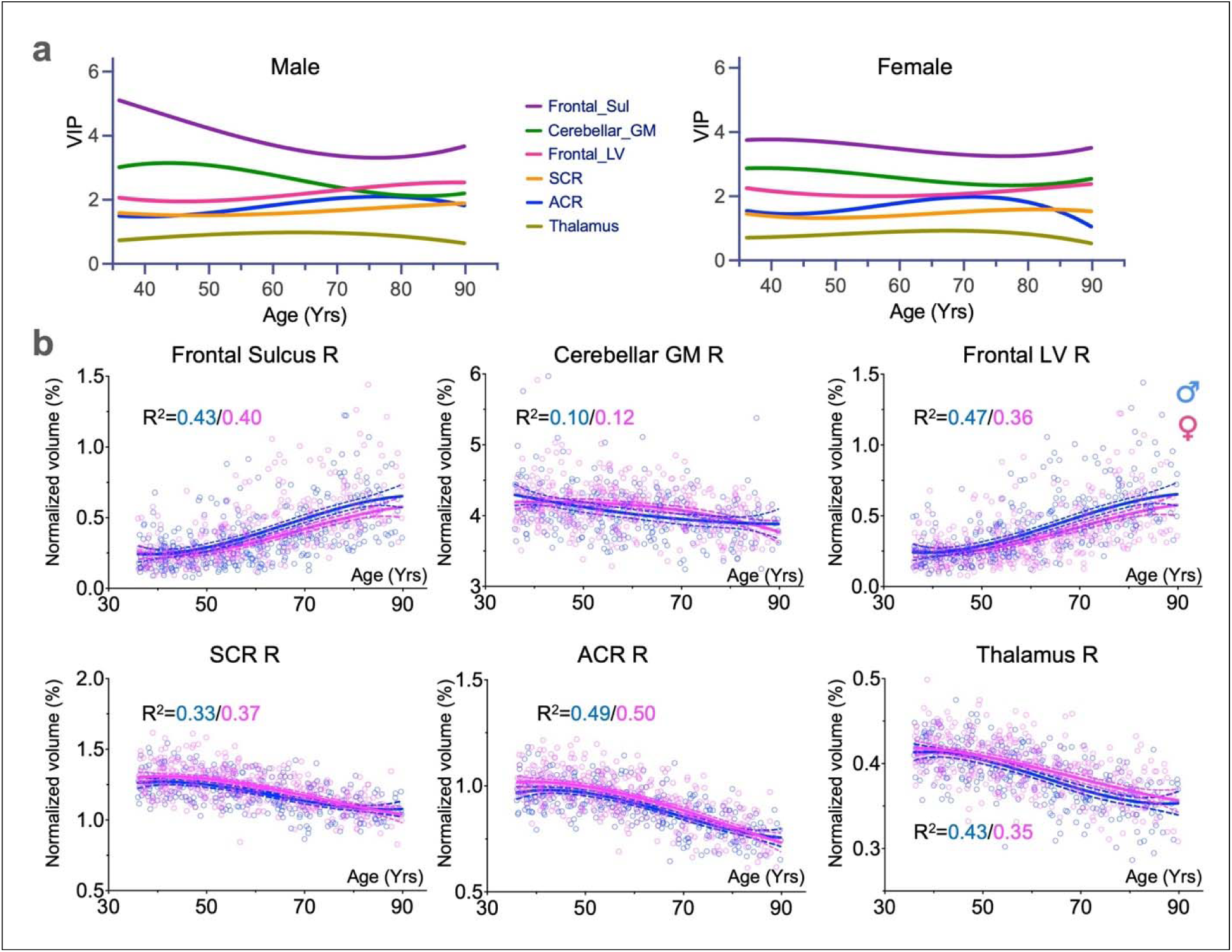
Age-dependent VIP scores in the HCP-A dataset. **a:** The age-related VIP score curves for selected brain regions in male and female HCP-A subjects. **b:** Scatter plots showing volume changes with age for six representative brain regions. Volumes of male (blue) and female (pink) subjects were plotted separately.

### Aging patterns on the manifold

The local aging patterns from individual neighborhoods can be aggregated to reveal broader trends. We applied K-means clustering on the coefficients of local PLSR of age, which were the regression weights and reflected how regional volume changes affect estimated brain age. We experimented with different numbers of clusters, finding that five clusters provided the most interpretable and stable results (**Fig. 5a**). The HCP-A subjects in each cluster occupied distinct and compact regions on the manifold (**Fig. 5b**), indicating that neighboring subjects shared similar broader aging patterns in regional volumes. Cluster 1 was associated with younger subjects, clusters 4 and 5 with older subjects, and clusters 2 and 3 with subjects spanning a wide age range.

**Figure 5:**
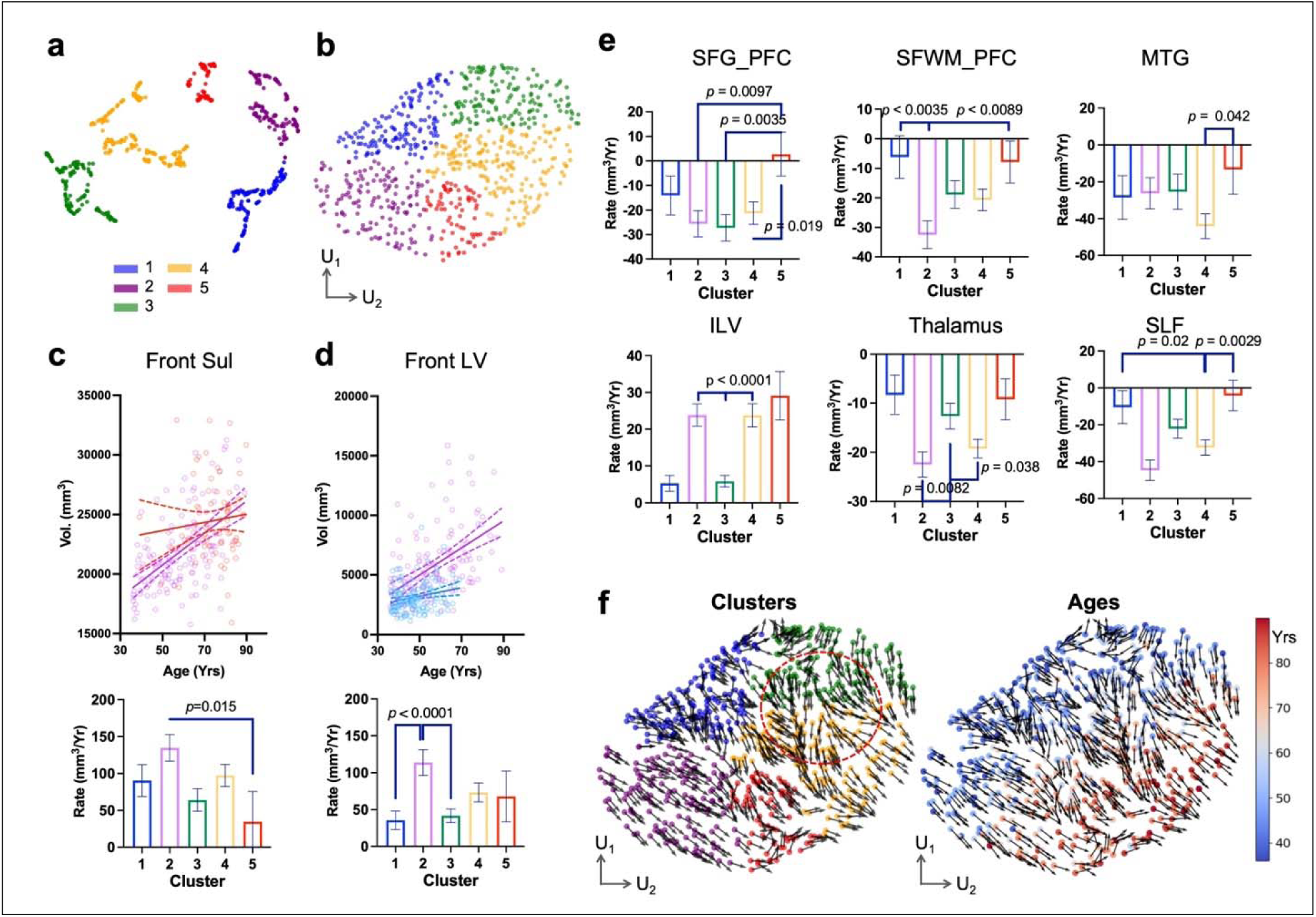
Distinct aging patterns on the manifold. a: Five clusters of HCP-A subjects based on locally estimated rates of volumetric changes in 72 brain structures with VIP scores greater than one. **b:** Distribution of the five clusters on the aging manifold. **c:** Frontal sulcus volume versus age for clusters 2 and 5, with linear regression fits showing estimated volumetric change rates (dashed lines indicate 95% confidence intervals). Cluster 2 exhibited significantly higher atrophy rates than cluster 5 (bottom panel). **d:** Frontal lateral ventricle volume versus age, demonstrating that cluster 2 showed significantly faster ventricular expansion than clusters 1 and 3. **e:** Comparison of volumetric change rates across key brain regions, revealing significant differences among the five clusters. **f:** Aging gradient vector field across the manifold with individual subjects colored by cluster assignment (left) and age (right).

For each cluster, we computed age-related volumetric change rates for high-VIP structures, exemplified by the frontal sulcus and frontal LV measurements (**Fig. 5c-d**). The five clusters exhibited distinct patterns of age-related volume changes. Cluster 1 (blue) and cluster 3 (green) showed moderate changes across most regions, though cluster 3 had notable changes in the prefrontal cortex (SFG_PFC). Cluster 5 (red) also exhibited moderate changes except in the inferior lateral ventricle (ILV). In contrast, clusters 2 and 4 demonstrated more pronounced regional specificity: cluster 2 (purple) showed accelerated frontal atrophy with faster ventricular expansion (frontal LV, inferior LV), sulcal widening, and atrophy of the superior frontal white matter (SFWM) and superior longitudinal fasciculus (SLF) (**Fig. 5e-f**). Cluster 4 (yellow) primarily showed changes in the MTG and inferior LV. Both clusters 2 and 4 showed faster thalamic atrophy than the other clusters. We also visualized aging gradients—volume changes corresponding to the greatest age increase—as vector fields across the manifold based on local PLSR results (**Fig. 5f**). Most aging vectors aligned with the age gradient (**Fig. 1d**), but a bifurcation occurred between clusters 3 and 4 (red circle in **Fig. 5f**), suggesting divergent aging pathways.

## Discussion

This study introduces a manifold-based framework for assessing individual brain aging based on regional volume measurements. This aging manifold provides a low-dimensional representation of regional volume configurations in the aging human brain, where age-related volumetric changes can be visualized as trajectories across the manifold (*22*). The approach enables computation of both overall brain age estimates and identification of key regional contributors that drive these aging patterns. The framework is adaptable to diverse MRI and non-MRI data.

Brain aging is inherently heterogeneous: no single region adequately captures its complexity (*32–36*). This challenge is further compounded by strong interregional correlations (*37–39*), which create redundancy in neuroimaging data and make dimension reduction essential for extracting latent variables that govern aging dynamics. Conventional methods such as PCA and independent component analysis have been previously applied (*13*, *34*, *38*, *40–42*), but their linear assumptions primarily capture global structure and may overlook subtle, region-specific (*27*) or nonlinear patterns across subgroups (*43*, *44*). Modern dimension reduction methods, including *t-*distributed stochastic neighbor embedding (*t*-SNE) (*45*) and UMAP (*26*), better preserve local topological relationships and can characterize nonlinear brain aging trajectories without linear constraints (*46*). Several groups have demonstrated the utility of these methods in uncovering local structures within MRI datasets (*22*, *23*, *47–50*). Building on this foundation, our aging manifold (**Fig. 1**) was derived from the HCP-A dataset, embedding high-dimensional regional volume measurements into 2D or 3D maps (**Supplementary Materials Figures S4 & 5**). For visualization clarity, we focused on the 2D manifold in this study, though the methodology can be readily extended to 3D or higher-dimensional representations as needed.

Unlike the distinct clusters often observed in gene expression classification or similar datasets (*51*), the manifolds derived from HCP-A structural MRI data as well as those reported previously (*22*, *23*) are relatively compact and continuous, with no apparent discrete clusters (**Fig. 1c**). This likely reflects the gradual nature of age-related brain changes and the multifactorial influences of genetic, environmental, and demographic variables on brain morphology (*52–54*). While HCP-A provides high quality MRI data from healthy aging participants, the manifold could be substantially improved by incorporating larger datasets (e.g. UK Biobank (*55*)) or harmonized multiple datasets (*18*, *19*). Such integration would (1) increase subject density in local manifold regions to improve inference accuracy, and (2) expand representation of under-sampled populations, particularly those over 90 years of age. A notable advantage of this framework is its ability to guide strategic data collection by identifying sparsely populated regions of the manifold. By doing so, it can predict the specific demographic and morphometric characteristics (e.g., brain volume, as shown in **Supplementary Materials Figure S4**) needed to optimize manifold coverage. Furthermore, incorporating pathological cases from datasets such as ADNI would extend the manifold beyond normal aging to include disease-related trajectories.

Although the manifold encodes topological rather than quantitative relationships, these relationships can be used for inference, as demonstrated by our brain age estimation analyses. Chronological age is the strongest known risk factor for late-onset AD (*56*, *57*), yet brain age more closely predicts disease likelihood (*58–60*). Deep learning-based brain age models, however, often suffer from limited interpretability, which hinders generalizability. To address this, methods such as layer-wise relevance propagation (*61*) and occlusion sensitivity analysis (*62*), have been used to identify regional contributions to brain age estimation. While informative, these methods are challenged by the high dimensionality of MRI data, including computational burden, multiple comparisons, spatial correlations, and artifacts from unrealistic occlusion. Manifold-based brain age estimation addresses many of these issues by operating in a low-dimensional space grounded in realistic aging data. Beyond providing estimates, the manifold offers additional advantages: by locating individual subjects within the manifold, we can assess extrapolation risk (e.g., proximity to sparsely sampled regions or manifold boundaries) and derive confidence intervals for predictions. This will ensure that inferences are grounded in biologically plausible aging patterns. It is important to note, however, that manifold representations are not unique – they depend on both the dimension reduction algorithm and the underlying dataset characteristics – highlighting the importance of optimizing manifold construction as a direction for future research.

Local analysis of individual neighborhoods on the manifold revealed region-specific volume changes that both confirm established findings and provide new insights into brain aging patterns. For example, the strong impact of cerebellar volume on brain age (**Fig. 3h**) is consistent with recent work by Raz et al. (*36*) and Bernard et al. (*63*), who demonstrated significant cerebellar atrophy in advancing aging – challenging earlier views that the cerebellum is largely spared during normal aging. Similarly, the pronounced contribution of frontal sulcus volume reflects frontal lobe atrophy (**Fig. 3e**), supporting the “last-in, first-out” hypothesis described by Fjell et al. (*1*, *64*), in which late-developing brain regions show earlier decline. Our observation of lateral ventricle enlargement (**Fig. 3g**) with age represents one of the frequently reported neuroimaging biomarkers of brain aging. White matter changes observed in this study provide further evidence of selective vulnerability: we found atrophy in the ACR and SCR, aligning with previous structural and diffusion MRI studies (*65*, *66*) that identified these projection fiber tracts as particularly vulnerable to age-related degeneration. Such vulnerability supports the disconnection hypothesis (*67*), which posits that age-related disruption of frontal-subcortical pathways contributes to cognitive decline, particularly in executive function and processing speed.

Our findings also align with prior data-driven reports that identified varying degrees of frontal, temporal, and subcortical atrophy as central markers for characterizing both heterogeneous normal aging and AD-related trajectories(*18*, *19*). Although the relatively small sample size of our normal aging dataset limited sensitivity to subtle aging patterns, the results nonetheless echoed earlier studies and revealed several interesting aging patterns. For example, cluster 2 (**Fig. 5**), characterized by marked frontal atrophy, included a higher proportion of male subjects – consistent with previous reports showing stronger frontal atrophy in males than in females (*68*). In contrast, clusters 3 and 4, colocalized with many of the female subjects, showed moderate and then more severe atrophy, especially in the temporal regions. This pattern may reflect hormonal changes associated with menopause, as demonstrated before (*69*).

One limitation of this study is its reliance solely on regional volume measurements for manifold construction, brain age estimation, and subsequent analyses. Incorporating additional MRI markers (e.g., cortical thickness, intensity, microstructure, and connectivity) or non-MRI biomarkers could likely enhance both the aging manifold’s representational capacity and brain age estimation accuracy (*8*, *70*). Interestingly, our attempts to integrate both structural and diffusion MRI yielded higher MAEs compared to using structural MRI alone. This unexpected finding may stem from residual mismatches between T_1_-weighted and diffusion MRI modalities, as well as partial volume effects that disproportionately impact diffusion metrics in complex tissue boundaries (*71*, *72*).

The local analysis assumes that structural and age similarities within a neighborhood are sufficient to establish homogeneity for aging analysis. While making the neighborhood smaller may better satisfy the homogeneity condition, this approach reduces the number of subjects per neighborhood and consequently diminishes the statistical power. Maintaining adequate statistical power requires large population datasets, yet subjects may still vary substantially in other critical factors—including genetics, health history, and lifestyle—that influence aging patterns.

In conclusion, this study introduces the aging manifold as a novel computational framework for evaluating individual brain aging. By enhancing interpretability, the methodology allows researchers to identify the specific factors that drive individual differences in brain aging patterns. This transparent framework advances our ability to model the complex, multidimensional nature of brain aging and provides insights into the heterogeneous pathways through which aging manifests.

## Materials and Methods

### Dataset

This study included 725 participants from the publicly available HCP-Aging database, Lifespan 2.0 release (https://www.humanconnectome.org/study/hcp-lifespan-aging/document/hcp-aging-20-release). Among the participants, 35 subjects were later excluded due to poor MRI data segmentation results as described later. The remaining 690 subjects ranged in age from 36 to 100 years (mean = 60.0 years, SD = 15.0 years) and was approximately balanced by sex (Male/Female =319/371). All participants were free of major neurological or psychiatric conditions (e.g., stroke, brain tumor, clinically diagnosed dementia). Detailed demographic information of HCP-Aging can be found in prior publications(*24*). The study protocol was approved by the institutional review board, and all participants provided written informed consent. The HCP-Aging data represents normal aging without known pathological conditions that affect cognitive abilities.

### Cognitive Function Assessment

Cognitive measures from the HCP-Aging dataset (https://www.humanconnectome.org/storage/app/media/documentation/LS2.0/LS_2.0_Release_Appendix_2.pdf) were incorporated into the UMAP embedding alongside regional brain volume features. The cognitive variables included Fluid Cognition Composite scores, Crystallized Cognition Composite scores, and Total Cognition Composite scores, all derived from the NIH Toolbox Cognition Battery(*73*, *74*).

Cognitive measures were obtained from the NIH Toolbox in the HCP-Aging dataset, including the Fluid, Crystallized, and Total Cognition Composites. The Fluid Composite assesses processing speed, working memory, episodic memory, and executive function, while the Crystallized Composite reflects premorbid intellectual abilities such as vocabulary knowledge and oral reading recognition. To capture both raw performance and age-normed scaling, we included six variables: nih_fluidcogcomp_unadjusted, nih_fluidcogcomp_np, nih_eccogcomp_unadjusted, nih_earlychildcogcomp_np, nih_totalcogcomp_unadjusted, and nih_totalcogcomp_np.

### MRI

We only used T_1_-weighted and multi-shell diffusion MRI data from the HCP-Aging dataset in this study. MRI data were acquired across four sites using Siemens Prisma 3 Tesla scanners equipped with 32-channel head coils, all running the same software version (E11C). A harmonized, state-of-the-art imaging protocol was implemented across sites, with standardized procedures, centralized training, and ongoing quality control analyses ensuring high inter-site consistency. T_1_-weighted MRI data was obtained using a magnetization-prepared rapid acquisition gradient echo sequence with the following parameters: voxel size = 0.8 × 0.8 × 0.8 mm³, echo times (TE) = 1.8/3.6/5.4/7.2 ms, repetition time (TR) = 2,500 ms, inversion time (TI) = 1,000 ms, flip angle = 8°, number of echoes = 4. Diffusion MRI data were acquired using a multiband 2D spin echo EPI sequence with the following parameters: TR/TE = 3,230 ms/89.20 ms, voxel size = 1.5 × 1.5 × 1.5 mm³, multiband acceleration factor = 4, b-values = 1,500 and 3,000 s/mm² (92∼93 directions per shell) with 28 non-diffusion-weighted (b = 0 s/mm²) images. Maps of fractional anisotropy (FA) and mean diffusivity (MD) were computed from diffusion MRI data using MRtrix 3.0 (www.mrtrix.org).

### Image segmentation and regional volumes

T_1_-weighted brain images were segmented using MRIcloud(*75*) into 286 regions according to the Johns Hopkins adult brain atlas(*76*). The quality of segmentation was visually inspected, and data from 35 subjects were excluded due to poor segmentation quality, and data from the remaining 690 subject were used in subsequent analysis. Among the 286 regions, the volumes of non-brain skull, optic tract, and bone marrow were excluded in this study. We also excluded the bilateral fimbria and rostral anterior cingulate white matter, which were not properly segmented in some subjects. We used the remaining 267 regional values for analysis. To align structural and diffusion data, T_1_-weighted images were registered to the mean diffusion-weighted image using the ANTs Symmetric Normalization (SyN) algorithm(*77*).

### UMAP

We computed the aging manifold using our in-house Python code built upon the open-source UMAP library (version 0.5.8; https://umap-learn.readthedocs.io)(26). A 267×690 matrix of subjects’ regional brain volumes was projected onto a low-dimensional manifolds using UMAP while preserving the local neighborhood structure of the high-dimensional volumetric data. The hyperparameters were tuned to favor settings that yielded an approximately uniform density of sample points within the boundary. After embedding a set of feature vectors in a low-dimensional space, UMAP also supports the online embedding of novel biomarker vectors from new subjects.

### Estimation of brain age and volume

A linear regression age prediction model was built using the low-dimensional manifold as the feature space. We used a distance-weighted kernel regression model with a kernel size (*k*) of 5, chosen to balance bias and variance: smaller *k*s produced unstable predictions due to noise, while larger *k*s overly smoothed local aging patterns. To evaluate the performance of age prediction, we employed a repeated leave-one-out strategy. In each run, *n* subjects were randomly selected to be placed on the manifold as samples, and the remainder *m* subjects were used for testing. The sample/test splits were stratified by age and sex, and random seeds were controlled for reproducibility. This procedure was run multiple times, and out-of-sample predictions were aggregated across runs to obtain final estimates. Model performance was quantified using mean absolute error and R². The same method was also applied to estimate regional volumes. All analyses were performed in Python 3.11 using the scikit-learn and matplotlib libraries.

### Partial least squares regression (PLSR)

PLSR projects regional volumes and chronological ages into respective latent spaces via linear transformations that maximize the covariance between the projected volumes and ages. For each subject on the manifold, we identified 50 nearest neighbors using the Ball Tree search. The neighborhood size from 25 to 100 were tested, and 50 was chosen to balance the need to avoid overfitting while preserving regional specificity. PLSR was performed with the number of latent variables set to 2∼5. The VIP score quantifies each region’s contribution to explaining variation in age, and regions with VIP scores above 1 were considered important. If a region’s VIP score exceeded 1 over the entire manifold, we classified it as a globally important feature.

### Clustering of the local PLSR results

The regression coefficients (βs) from local PLSR quantify regional volumes’ association with age. Clustering of local βs on the manifold was performed using UMAP with the following parameters: n_components = 2, n_neighbors = 15, min_dist = 0.1, and random_state = 2. We then applied k-means clustering (n_clusters = 3∼6, random_state = 30) to partition the embedding into discrete groups, and n_clusters = 5 produced more robust than others.

### Generation of the aging gradient vector field

Within each neighborhood on the manifold, we performed linear regression to find a linear approximation of chronological age as a function of the two UMAP coordinates. The local gradient vectors from the regression indicated the direction of volume changes that corresponded to the fastest aging. These gradient vectors were normalized to the same length to visualize directions rather than magnitudes, which may not be directly comparable on different parts of the manifold.

## Author contributions

Conceptualization: ZL, CJS, JZ; Methodology: ZL, CJS, CL, JL, JZ; Investigation: ZL, CJS, CL, JL, JZ, HR, YG; Visualization: ZL, CJS, JZ; Supervision: JZ; Writing—original draft: ZL, CJS, JZ; Writing—review & editing: ZL, CJS, JZ, HR, YG

## Data and materials availability

All data needed to evaluate the conclusions in the paper are available from HCP and ADNI websites.

