## supplemental materials for "A Manifold Framework for Interpretable Brain Age Estimation and Aging Trajectory Mapping"

Zifei Liang *et al.*

*Jiangyang Zhang.

**This PDF file includes:**

Figs. S1 to S8

Tables S1

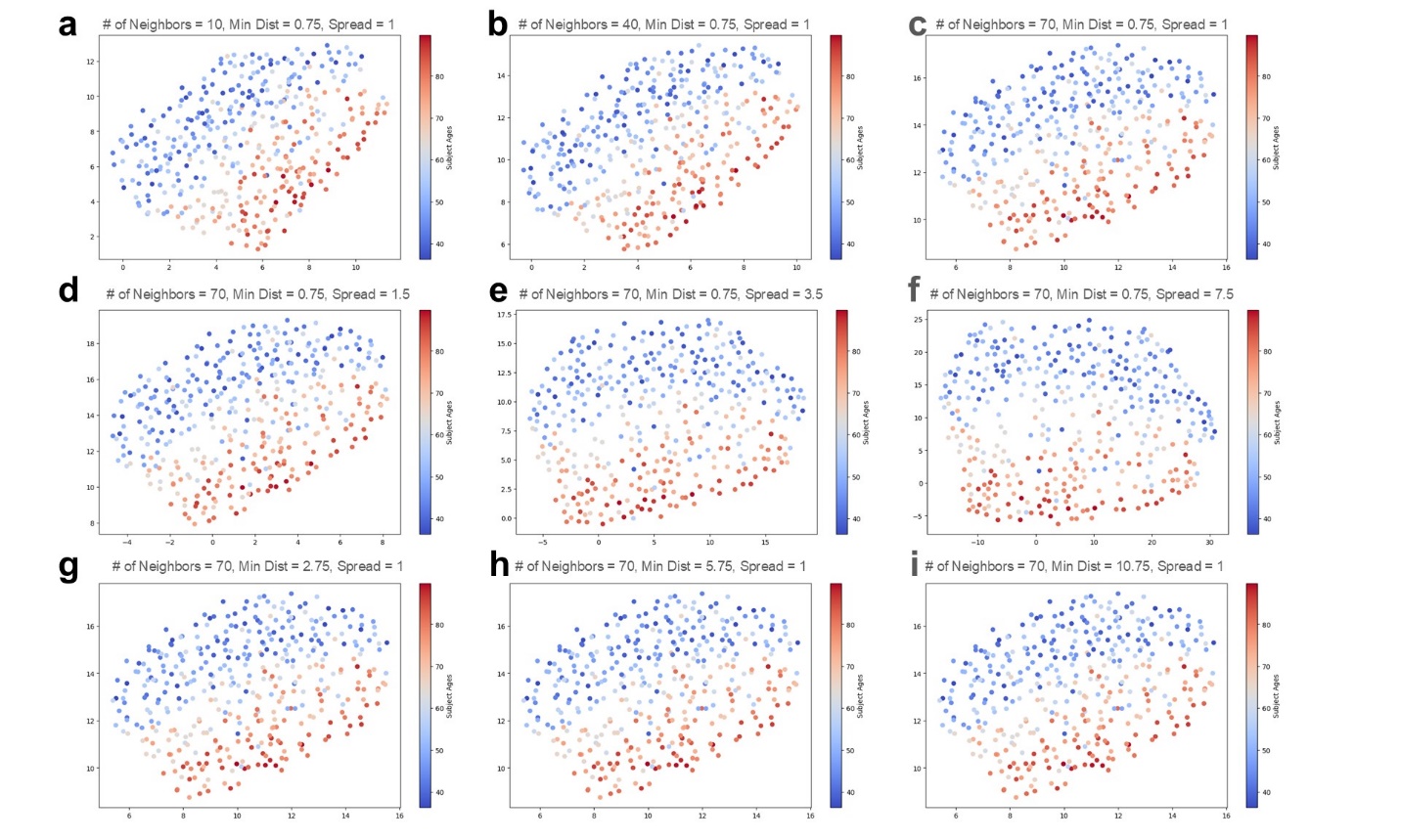

**Figure S1:** The manifolds generated by UMAP from HCP-A regional volume data under different settings by varying three main parameters: number of neighbors (# of Neighbors), minimum distance (Min Dist), and spread. The color indicates subject’s age.

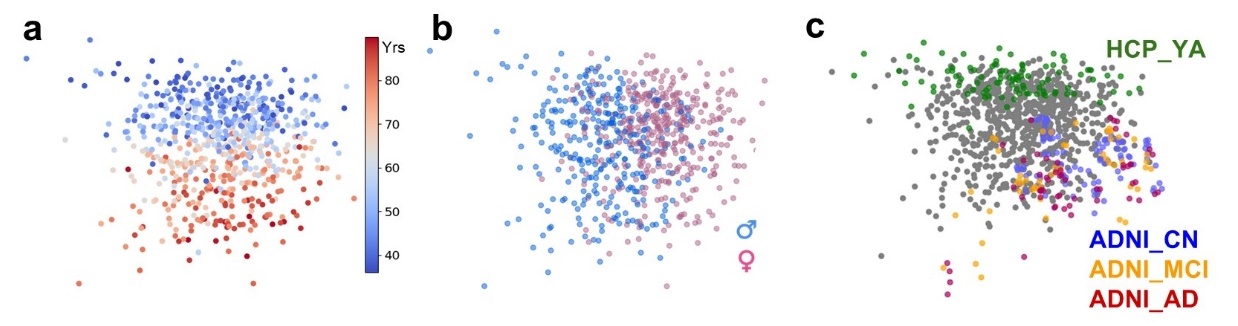

**Figure S2:** Dimensional reduction of the HCP-A structural MRI data using PCA. **a-b:** Distribution of HCP-A subject data along the largest two eigen-vectors, with points color-coded by age (**a**) and sex (**b**). **c:** Distribution of the HCP-YA and ADNI data in **Figure 1g** projected into the PCA eigenspace.

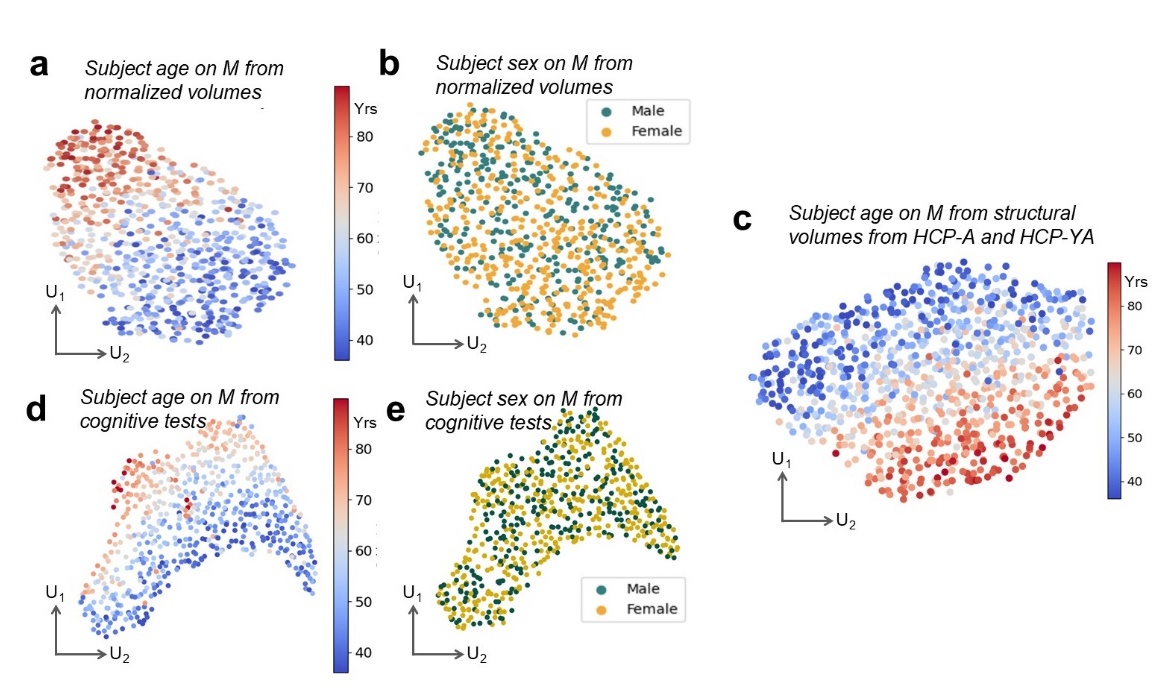

**Figure S3:** The aging manifold can accommodate new data and non-MRI data. **a-b:** The manifold based on HCP-A regional volume data normalized by the whole brain volume and the distributions of subject age and sex. **c:** The distribution of subject age on a manifold based on regional volumes of 690 HCP-A data and 200 randomly selected HCP-YA data (20-35 years old, F/M=100/100). We chose 200 subjects to balance the number of subjects in each decade. **d-e:** The distribution of subject age and sex on a manifold based on the cognitive test results (unadjusted by age) from the HCP-A dataset.

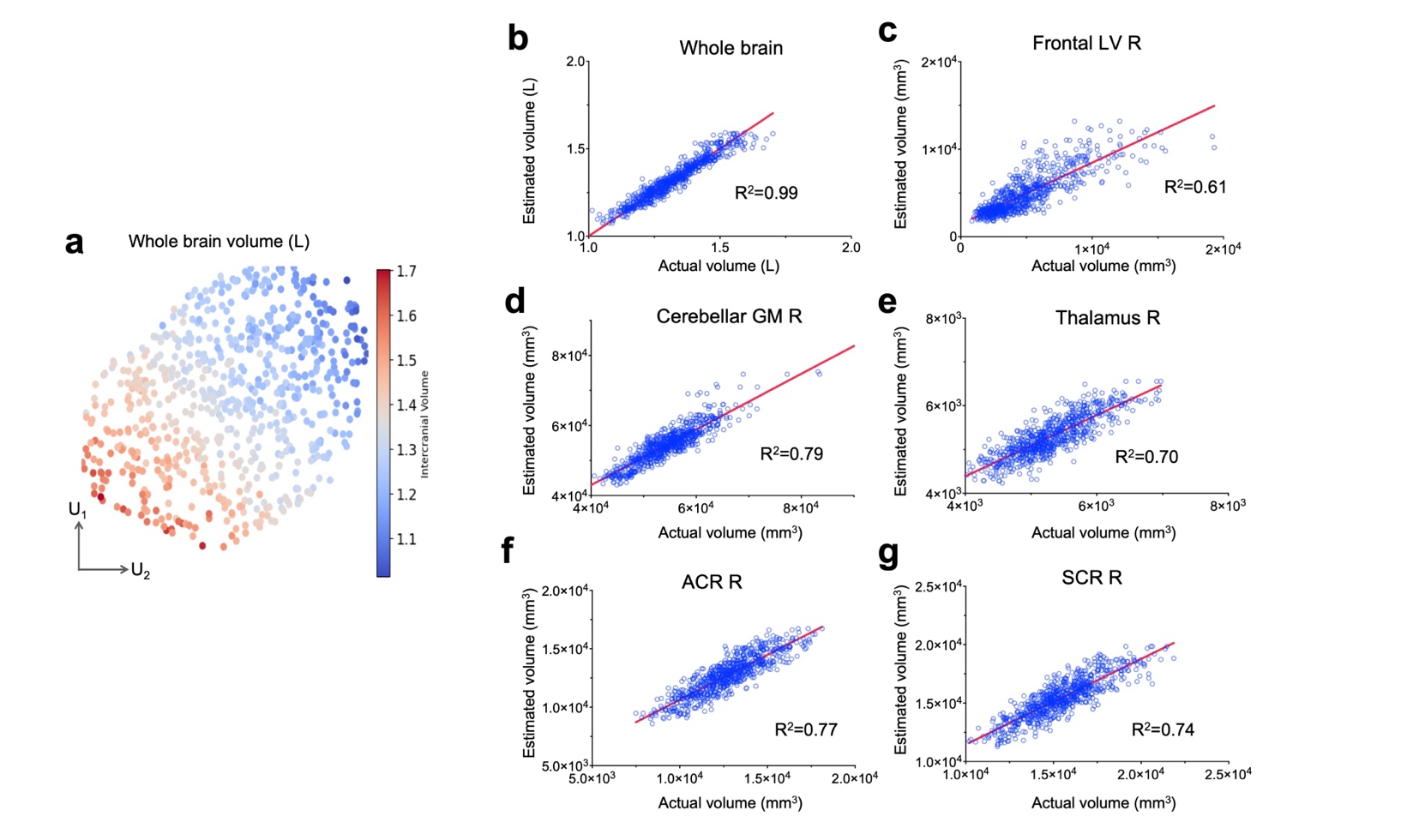

**Figure S4:** Regional volume on the manifold and estimation of regional volumes by regression. **a:** Distribution of whole brain volumes of HCP-A subjects on the manifold. **b:** The correlation between estimated whole brain volumes using local regression and the actual whole brain volumes. **c-g:** correlations between estimated and actual and right frontal lateral ventricle (Frontal LV R), right cerebellar gray matter (Cerebellar GM R), right thalamus (Thalamus R), right anterior corona radiata (ACR R), and right superior corona radiata (SCR R) volumes.

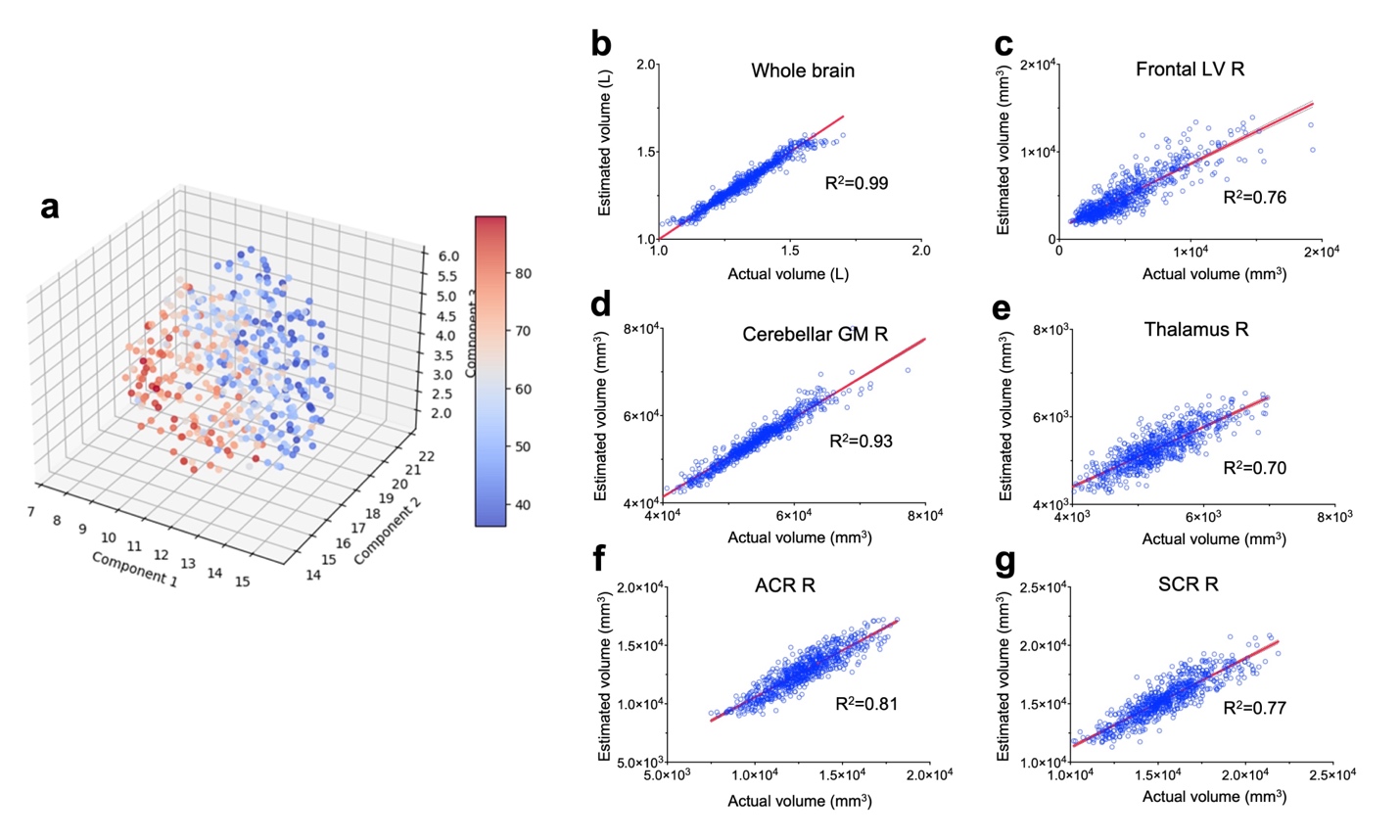

**Figure S5:** Regional volumes on a 3D manifold and estimation of regional volumes by regression. **a:** Distribution of age of HCP-A subjects on the 3D manifold. **b:** The correlation between estimated whole brain volumes using local regression and the actual whole brain volumes. **c-g:** correlations between estimated and actual and right frontal lateral ventricle (Frontal LV R), right cerebellar gray matter (Cerebellar GM R), right thalamus (Thalamus R), right anterior corona radiata (ACR R), and right superior corona radiata (SCR R) volumes.

**Figure S6:** Volume correlation matrix among the structures in the HCP-A dataset.

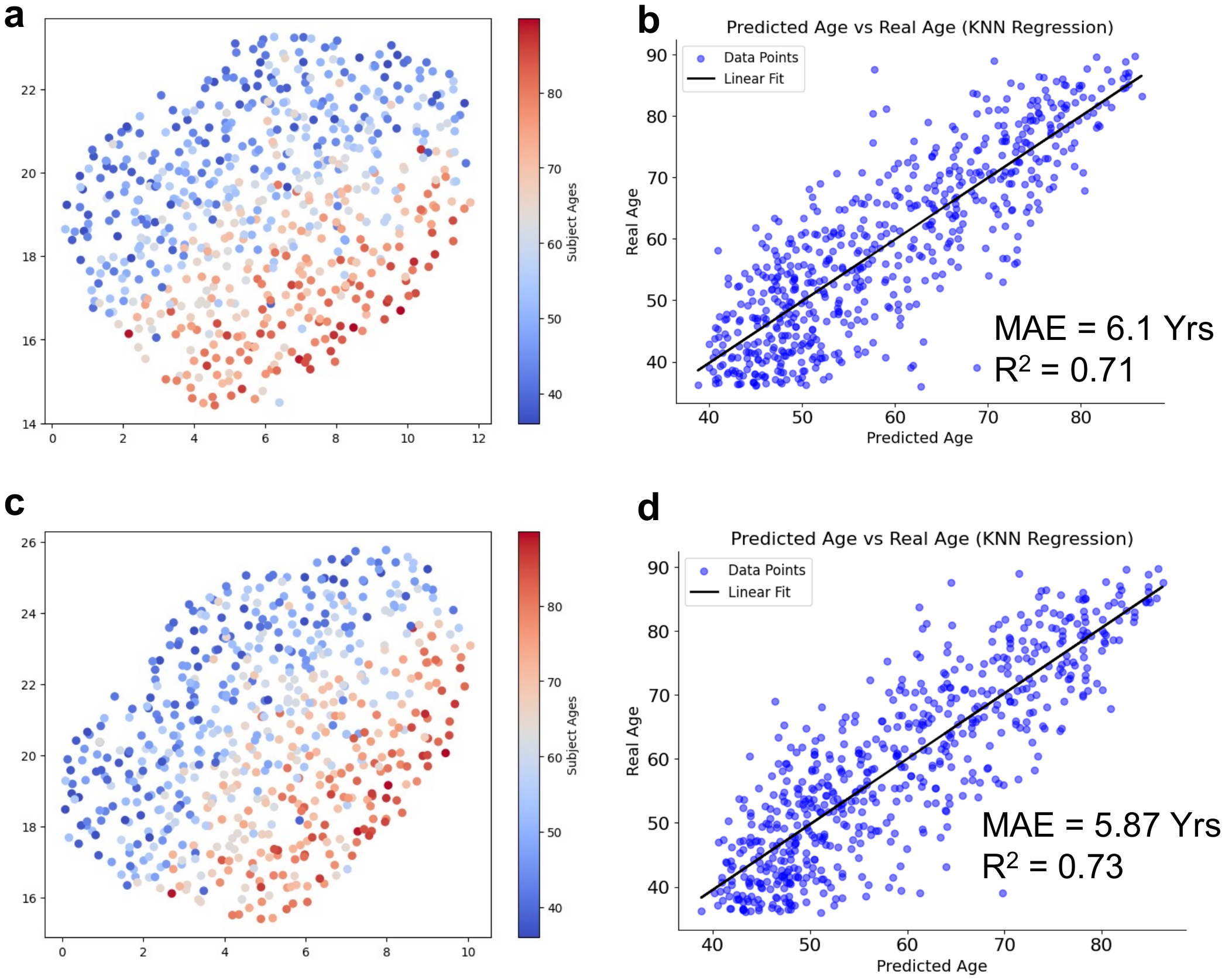

**Figure S7:** Modified brain aging manifolds based on the PLSR results. **a-b:** A manifold based on the volumes of 72 brain structures with VIP scores greater than 1 and the correlation plot between estimated and chronological ages of HCP-A subjects. **c-d:** The manifold based on the 72 regional volumes plus cognitive measurements from HCP-A data and the corresponding correlation plot.

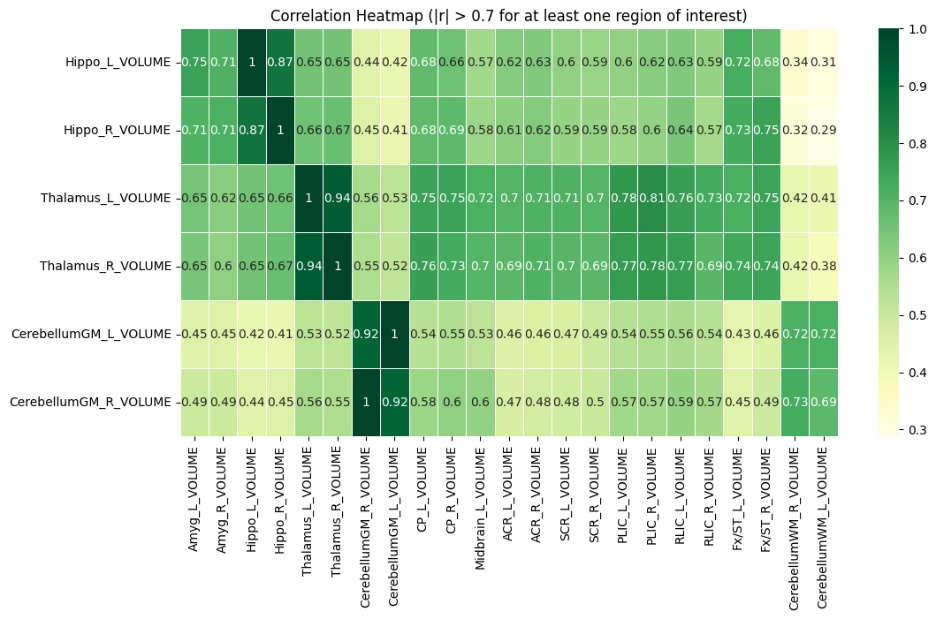

**Figure S8:** Correlations among selected structures in the HCP-A dataset.

**Supplementary Table S1:** Abbreviations of the 32 brain regions with VIP scores greater than 1.

| **Abbreviation** | **Full name** |
| --- | --- |
| FrontSul | sulci of the frontal lobe |
| ParietSul | sulci of the parietal lobe |
| SylTempSul | Sylvian fissure and posterior insular sulcus |
| OcciptSul | sulci of the occipital lobe |
| CentralSul | Central sulcus |
| LV_Frontal | frontal horn of the lateral ventricle |
| LV_body | body of the lateral ventricle |
| LV_atrium | Lateral ventricle, atrium part |
| PrCWM | subcortical white matter of the precentral gyrus |
| ACR | Anterior corona radiata |
| PoCWM | subcortical white matter of the postcentral gyrus |
| SCR | Superior corona radiata |
| SLF | Superior longitudinal fasciculus |
| LWM | subcortical white matter of the lingual gyrus |
| SCC | Splenium of corpus callosum |
| SFWM_PFC | subcortical white matter of the superior frontal gyrus/ prefrontal cortex |
| MOWM | subcortical white matter of the middle occiptial gyrus |
| MTWM | subcortical white matter of the middle temporal gyrus |
| PCC | Posterior cingulate cortex and subcortical white matter |
| dorsal_ACC | dorsal Anterior cingulate cortex and subcortical white matter |
| CerebellumGM | cerebellum gray matter |
| STG | superior temporal gyrus |
| MFG_DPFC | Middle frontal gyrus (dorsolateral prefrontal cortex) |
| SFG | Superior frontal gyrus |
| MTG | Middle temporal gyrus |
| MOG | middle occipital gyrus |
| AG | angular gyrus |
| PrCG | Precentral gyrus |
| SFG_PFC | superior frontal gyrus/ prefrontal cortex |
| MFG | Middle frontal gyrus |
| SMG | Supramarginal Gyrus |
| FuG | Fusiform gyrus |
| SPG | SUPERIOR PARIETAL LOBULE |
| PoCG | Postcentral gyrus |
| ITG | inferior temporal gyrus |
